# Elusive scents: neurocomputational mechanisms of verbal omissions in free odor naming

**DOI:** 10.1101/2025.02.13.637907

**Authors:** N. Chrysanthidis, R. Raj, T. Hörberg, R. Lindroos, A. Lansner, E.J. Laukka, J.K. Olofsson, P. Herman

**Author notes:** corresponding author: Pawel Herman.

## Abstract

Odor naming is considered a particularly challenging cognitive test, but the underlying cause of this difficulty is unknown. People often fail to report any source label to identify common odors, resulting in omissions (i.e., a lack of response). Here, with the support of a computational model, we offer a hypothesis about the neural network mechanisms underlying odor naming omissions. Based on an evaluation of behavioral data from almost 40,000 odor naming attempts, we suggest that high omission rates are driven by odors that are referred to by multiple linguistic labels. To explain this observation at the systems level, where olfactory perception and language (semantic) processing are produced by interacting cortical systems, we developed a computational model consisting of two associatively coupled attractor memory networks (odor and language networks), and investigated the effect of Hebbian-like learning on the simulated task performance. We used distributed network representations for the odor percepts and word label mental objects, and accounted for their statistical inter-relationships (correlations) extracted from collected data on odor perceptual similarity, and from a large Swedish odor language corpus, respectively. We evaluated a novel hypothesis, that Bayesian-Hebbian synaptic plasticity mechanisms can explain behavioral omissions in odor naming tasks, casting new light on the underlying mechanisms of this frequently observed memory phenomenon. Due to the nature of Bayesian-Hebbian associative learning connecting the two networks, there was a progressively weaker coupling for odors paired with multiple different labels in the encoding process (one-to-many mapping). Thus, when the model was cued with perceptual odor stimuli that established multiple word label associations (one-to-many mapping), the olfactory language network often produced subthreshold network responses, resulting in elevated omissions (opposite to one-to-few mapping scenario that led to improved performance scores). Our results are of theoretical interest, as they suggest a biologically plausible mechanism to explain a common, but poorly understood, behavioral phenomenon.

## Introduction

Multiple lines of experimental evidence show that odors are difficult to verbally name even if the odors are familiar (Engen and Pfaffman, 1960; Desor and Beauchamp, 1974; Cain, 1979; de Wijk and Cain, 1994). Naming perceptual objects constitutes a complex cognitive process that involves distributed language networks in the brain (Olofsson and Gottfried, 2015; Binder et al., 2009). Yet, naming odors (i.e. verbalizing the prototypical source objects from which they originate, e.g. “lemon” or “gasoline”) is less consistent and less accurate when compared to visual or auditory stimuli. This has been partly attributed to a so-called “weak link” between odors and words (Herz and Engen, 1996). There are different hypotheses about the underlying mechanisms of this suggested weak association between odor and language processing pathways in the brain, either pointing to the structural organization and functional connectivity between the relevant neural network resources in the human cortex as the main problem (Olofsson and Gottfried, 2015; Olofsson et al., 2013; Olofsson et al., 2014), or implying lexical deficiencies in the olfactory vocabulary of some languages (Majid and Burenhult, 2014; Majid, 2015). The exact behavioral manifestations of this presumed intrinsic difficulty in odor naming depend on the experimental paradigm but can be generally categorized as either mistakes (incorrect naming) or omissions (lack of any verbal response to an odor stimulus). Considerably less attention has been paid to the latter phenomenon and, consequently, there have been hardly any direct hypotheses about the mechanistic origins of such “blank” responses (omissions). Although omissions have been observed for several familiar odors (Hörberg et al., 2024), it remains unclear how odor-specific the omissions are, and there is currently no computational model that may explain the phenomenon.

### Causes of odor naming omissions

Evidence from studies on brain lesions and neurocognitive experiments demonstrate that damage to the areas which encode odor representations or their potential learned semantic, lexical and episodic associations (Gottfried et al., 2004; Cerf-Ducastel & Murphy, 2006; Olofsson et al., 2013, 2014; Seubert et al., 2020) could impair the associative pathways and lead to loss of odor associations, even if the fundamental sensory information may remain intact (Eichenbaum, Morton, Potter, & Corkin, 1983). Brain damage or natural aging processes also lead to olfactory memory loss (Josefsson et al., 2017; Olofsson et al., 2016, Devanand, 2016; Ekström et al., 2017; 2020; Stanciu et al., 2014). However, loss of odor episodic associations (Tulving, 1972, Duff et al., 2020; Habermas et al., 2013; Viard et al., 2007) may emerge at faster time scales (compared to, for example, aging effects), and depend on memory overload, i.e., some odor representations can be linked with an excessive number of different episodic contexts or share many other competing memory associations (one-to-many mapping) (Opitz, 2010). Behavioral studies demonstrated that visual perceptual objects can be rapidly decoupled from their multiple memory associations, which in the realm of episodic memory is sometimes referred to as decontextualisation (Opitz, 2010). The multiplicity of memory associations has been demonstrated to result in the decrease of episodic memory recall performance (Opitz, 2010; Smith and Manzano, 2010; Smith and Handy, 2014). Baddeley (1988) hypothesized that semantic memory may reflect the extraction of information over multiple learning episodes (information detached from episodic details). Given the rather significant linguistic diversity in naming odors (i.e., multiple odor-label associations learned for the same odors over a lifetime, see, e.g. Majid et al., 2018; Lehrner and Walla, 2002), this hypothesis of memory overload (one-to-many mapping) might also hold true for odors and their associated correlates.

### Language diversity in odor naming tasks

Iatropoulos et al. (2018) demonstrated that some odor labels (e.g. “heavy”) may be applied to a large set of odors, thereby increasing the number of odor-label associations, and this can be viewed as a higher memory load associated with those odors. In that study, novel metrics were developed to characterize the semantic content of labels related to olfaction. They found that some labels are used selectively for certain odors (high specificity, e.g. “cinnamon”) but with lower frequency while others are more olfaction-unspecific, yet used frequently to describe a large family of odors that possibly share common features (low specificity), e.g., citrus fruits. In other words, even though some odors could be uniquely characterized by odor-specific labels, they can still be referred to by a broader class of labels (less odor-specific) that are thus used more frequently (e.g., “sour”). Furthermore, Majid and Burenhult (2014) showed that English-speaking subjects provided less unique word labels to name odor stimuli compared to naming color cues. Thus, multiple lines of evidence have demonstrated the variability of linguistic labels used in odor naming.

In this work, we first examined the hypothesis that the language variability and diversity in selecting word labels to refer to odors may explain the frequent phenomenon of verbal omissions reported in free odor naming scenarios. To this end, we analyzed experimental data collected in the Swedish National Study on Aging and Care in Kungsholmen (SNAC-K), where participants underwent an odor naming test and were asked to freely name and then identify the presented odors using multiple-choice cues (Larsson et al., 2016). Results suggest that the odors referred to by many different labels by different participants (high variability among participants) are also characterized by an elevated rate of verbal omissions. To gain insights into neurocomputational mechanisms related to this observation, we built a computational attractor memory model and proposed a synaptic plasticity basis as the key factor underlying the observed trend for verbal omissions in odor naming. In particular, we examined a synaptic learning hypothesis related to effects of language variability in omissions using a Bayesian-Hebbian learning rule implementing long-term memory associations between odor percepts and the corresponding word labels. We then simulated a free odor naming test on the subject population level in line with an experimental paradigm used in the SNAC-K study (a single model accounting for collective statistical characteristics of the odor naming performance of the SNAC-K subjects) and evaluated the average quantitative outcomes across the palette of available perceptual odor objects. In addition, taking advantage of the model we also analyzed other factors that may contribute to omissions, such as cross-talk among distributed neural representations of both odor percepts and word labels encoded in the network model, as well as bi-directional interactions between the corresponding olfactory and language systems. Importantly, our model provides deeper mechanistic insights and predictions derived from a framework that is developed to explain the origins of high omission rates in free odor naming tasks. The overarching aim of this study is to acquire a better fundamental understanding of the neuronal and synaptic plasticity processes underlying behaviourally quantifiable odor naming effects.

## Results

### The omission rates and the diversity of labels used for odor naming

Our original hypothesis about the origins of omissions in free odor naming involved the diversity of word label associations with a given odor. In particular, we proposed that associating an odor perceptual representation with a diverse set of word labels in the long-term process of encoding episodic memories throughout the subject’s life increases the risk of omission when the subject is exposed to that odor stimulus and is requested to name it. To validate this proposition we first analyzed SNAC-K experimental data (see Methods - Behavioral data), which enabled us to investigate this effect on the average population rather than individual subject level. In other words, we established that the diversity of labels used collectively by the study participants to name the odors correlated with the omission rates. We quantified the diversity of odor labels using Simpson’s diversity index, which takes into account both the type and frequency of word labels per odor (see *Methods - Characterizing odor-label diversity*). We observed indeed a significant correlation between high omission and high heterogeneity of odor labels; that is, odors that were referred to by several labels (0 Simpson index indicates high label diversity) resulted in elevated omissions (Fig. 1, linear regression, r=-0.46, p<0.05).

**Figure 1:**
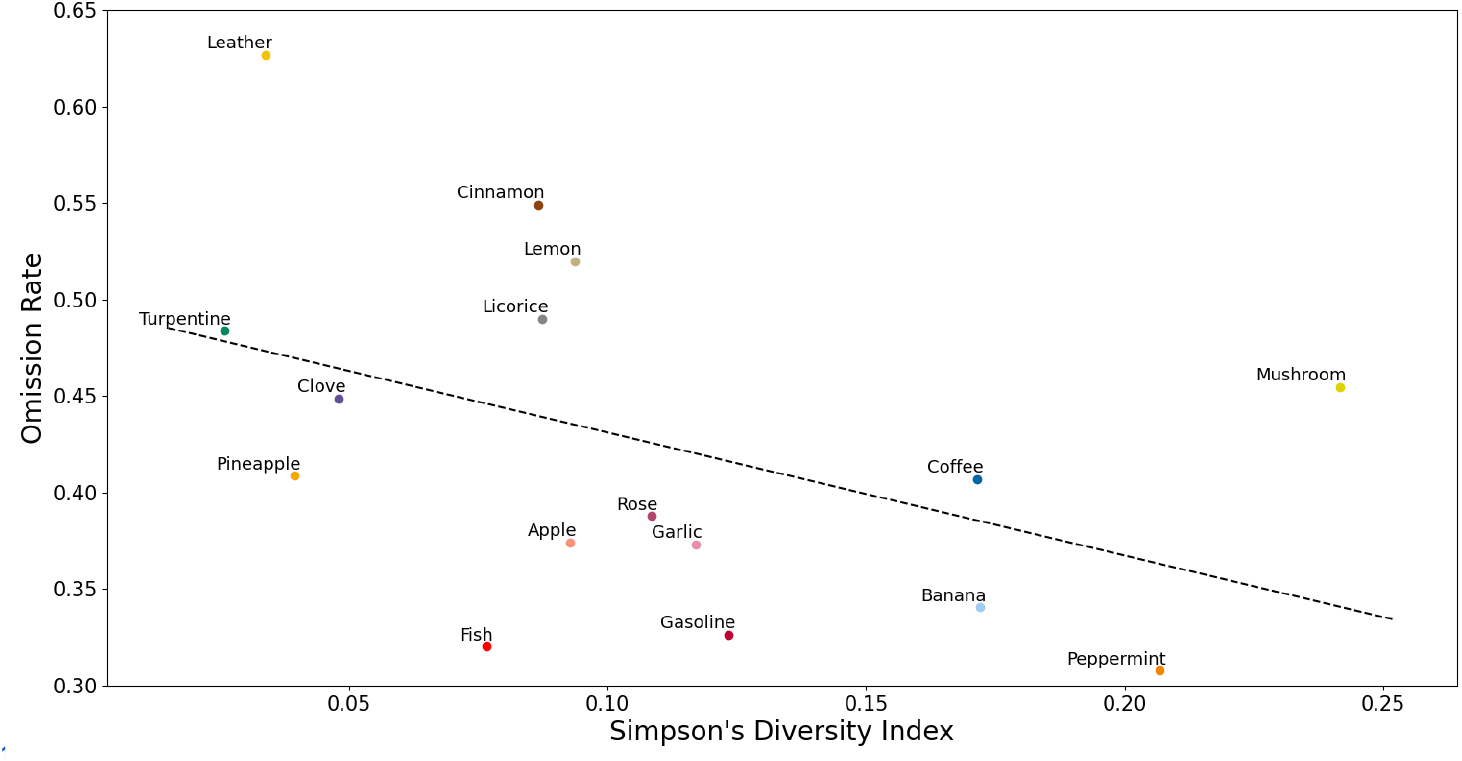
Language diversity index for all the SNAC-K odors. For each odor stimulus we calculated the label diversity using the Simpson Diversity Index (see *Methods - Characterizing odor-label diversity*) and the omission rate. There is a significant negative correlation between the two quantities (r=-0.46, p<0.05, n = 16).

### Odor-label diversity and omissions in the model

To explore potential neurocomputational processes underlying the effect of verbal omissions in free odor naming, we built a neural network model composed of two networks, the so-called odor and language networks (see *Methods - Network model*). At first, we represented odor percept and word-label objects (memory patterns) as distributed neural codes reflecting the perceptual and semantic, respectively, pairwise similarities derived from the available data (Lindroos et al., 2022; see details in *Methods - Odor percept and label representations derived from data*). These objects, corresponding to odor percepts and word labels, were then pre-encoded independently in their respective networks by means of the Bayesian-Hebbian learning rule (Sandberg et al., 2002; see *Methods - Plasticity learning rule*). In particular, multiple epochs of sequential training with a slow learning rate were applied to each network to account for the long-term consolidated nature of these memory objects (see details in Methods - Memory pre-encoding).

Before we simulated a free odor-naming experiment inspired by the SNAC-K study, we had to bind the pre-encoded word label representations in the language network with the odor percept representations in the odor network. These between-network connections (the synaptic weights and neural biases, see *Methods - Memory pre-encoding*) were trained with the aforementioned long-term Bayesian-Hebbian learning in a paired cue paradigm, i.e. we simultaneously stimulated the language and odor memory networks with the matching cues in the spirit of Hebbian co-activation (Sandberg et al., 2002). This process of matching odor percepts with word labels was meant to reflect the human declarative memory system where perceptual experiences are coupled with object names via paired-associate learning (in this case, coupling of odors with odor names). To examine our hypothesis about the relationship between omission rate when attempting to name an odor and the diversity of word labels associated with that odor, we matched the training cues during the simulated process of paired-associate learning based on the statistics derived from the behavioral responses reported in the free odor naming part of the SNAC-K study (Larsson et al., 2016). In particular, building on our previous analysis for each odor, we identified a set of word labels most frequently used to name it (irrespectively whether it was correct or not) collectively by the SNAC-K study participants. The number of word mappings for each odor percept was determined based on the Simpson’s diversity index (Majd et al., 2018; see *Methods - Characterizing odor-label diversity*). In particular, we related four modes in the histogram of the Simpson’s diversity index to the number of mappings ranging from 1 to 4 (four distinct families of odors were identified). This resulted in the odor-label associations shown in Fig. 2A. The odors-label associations were decided by the frequency of descriptor occurrences in the SNAC-K data (responses from participants), for example, odors forming four associations were associated with the top four most frequent descriptors in the behavioral data.

**Figure 2:**
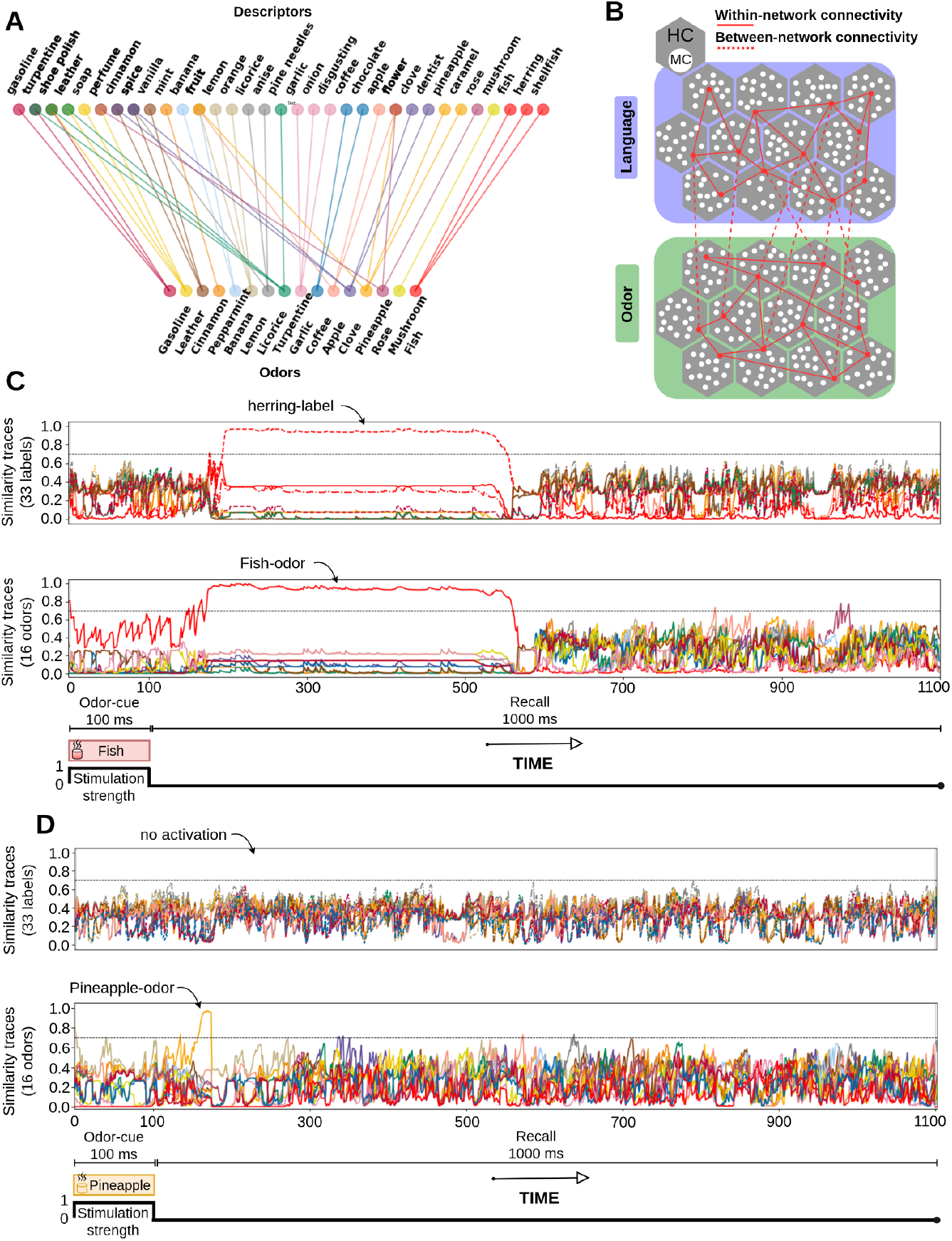
Pre-encoding of odor and label associations in a two-network memory model. **A**) Four distinct families of odors were identified based on their diversity index and omission rates (cf. Fig. 1A). Therefore, the odor groups reflected the number of odor associations with different labels (from one-to-one [minimum] to one-to-four mappings [maximum]). These odor-label associations were trained and preloaded in the two network system representing long-term consolidated memories (see Methods for details). **B**) Schematic of the odor (green) and language (blue) networks. Within-network connectivity reflects synaptic connections, considered to result from long-term memory consolidation, between units distributed across the modules (hypercolumns) of the same network that constitute neural ensembles - memory representations. Between-network connectivity is formed by associative connections between units of the two networks, established due to episodic memory binding processes. The odor network stores 16 odor percept memory objects and the language network stores 33 word labels. Their representations were extracted to reflect statistical similarity relationships estimated from the available data (see Methods). **C**) Trial structure of the simulated task: the odor network is stimulated for 100 ms with a partial cue (see *Methods - Partial pattern activation during odor naming task in the model*) corresponding to a selected odor percept with the following recall phase of 1000 ms (bottom). During the recall phase, we analyze the response activity in the language network that serves as the “verbal” readout. Responses of the odor (middle) and language (top) networks are quantified with the traces of similarity between the evoked activity patterns and the pre-encoded odor precepts and word labels, respectively. In that particular instance, the Gasoline odor percept was cued for 100 ms (brief cues are sufficient to trigger the model’s response to odor stimuli). The threshold for the network’s readout is marked with a dotted line. Label-tags shown in the plot correspond to either a cue identity (“Fish” for the odor network) or the word label output (readout for the network’s suprathreshold response) of the language network (“fish” and “herring”). Successful activation (above threshold activity) of the odor gasoline leads to correct retrieval of one of the associated word-labels in the language network (“herring”). The use of capitalized descriptors for odor percepts, e.g. “Fish”, is on purpose to distinguish from word labels, e.g. “herring”. **D**) Omission in the language network. The cued odor percept “Pineapple” is activated in the odor network, yet the language network remains silent (subthreshold activity), thus the odor activation does not lead to a retrieval of an associated word label descriptor.

Following the process of paired-associate learning of network memory representations of the matched odor percepts and word labels we simulated the free odor naming task in SNAC-K study. To this end, we cued each of the 16 stored odor perceptual memories for 100 ms (cued odor network, Fig. 2B) and in the subsequent recall block of 1 s we recorded the responses of both networks. Each network’s response was quantified using a measure of similarity (cosine similarity, see *Methods*) between the distributed pattern of network activation and the memory representations of all stored objects. This approach allowed us to track the odor percept and word label network responses over time. Once the resulting convergent state was sufficiently similar to one of the stored memory states (i.e. the recall detection threshold *r*_*th*_ was exceeded for at least a certain duration 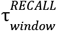, see Table 1 in *Methods*), that memory activation (i.e. the odor percept for the olfactory network or word label for the language network) was considered as a perceptual or verbal output, respectively. The subthreshold activation of the language network within the 1-s-long recall block (i.e. no legitimate output) was considered as a verbal omission. A subthreshold response could be a result of either the network’s transient activity (without convergence) or the network’s convergence to one of so-called spurious attractor states, i.e. “false” memory states that do not coincide with any of the pre-encoded memories. In Fig. 2C we show an instance of a cued odor percept activation corresponding to “Fish” that leads to correct retrieval of its word label “herring” in the language network after a short delay. We also show an example of a verbal omission in Fig. 2D where the stimulation of the odor percept “Pineapple” does not lead to an activation in the language network (blank response), even though the odor “Pineapple” was briefly activated in the odor network.

**Table 1:** Model and synaptic parameters.

| Model parameter | Symbol | Value | Plasticity parameter | Symbol | Value |
| --- | --- | --- | --- | --- | --- |
| Network size | HCs x MCs | 15 x 15 | BCPNN gain (within network) | $g_w$ | 1.6 |
| Number of hypercolumns | HC | 15 | BCPNN gain (between networks) | $g_w^{asso}$ | 0.4 |
| Number of networks | $N_{net}$ | 2 | BCPNN bias gain | $g_b$ | 1 |
| Number of minicolumns | MC | 15 | BCPNN lowest probability | $\epsilon$ | 0.01 |
| Adaptation time constant | $\tau_w$ | 0.01 s | P trace time constant | $\tau_p$ | 20 s |
| Adaptation gain | b | 0.28 | Z trace time constant | $\tau_z$ | 1 ms |
| Recall detection threshold | $r_{th}$ | 0.3 | Plasticity - encoding | $\kappa_{encoding}$ | 1 |
| Stimulation | $r_{STIM}$ | 100 ms | Plasticity - recall | $\kappa_{recall}$ | 0 |
| Recall threshold | $r_{RECALL}$ | 1000 ms | Connection probability | $cp_{within-net}$ | 1 |
| Recall time window | $\tau_{window}^{RECALL}$ | 40 ms | Connection probability | $cp_{between-net}$ | 0.8 |

HC: Hypercolumn, MC: Minicolumn

Having simulated recall for all available odors in the model, we found evidence that odors associated with many different labels in the prior odor-label learning (one-to-many mapping, Fig. 2A) tended to generate more omissions, i.e. blank responses, in the language network (Fig. 3A, Model). This tendency is in line with the correlation effect reported in Fig. 1 and, consequently, matches the behavioral omission rates in the SNAC-K study for the corresponding groups of odors (Fig. 3A, SNAC-K data).

**Figure 3:**
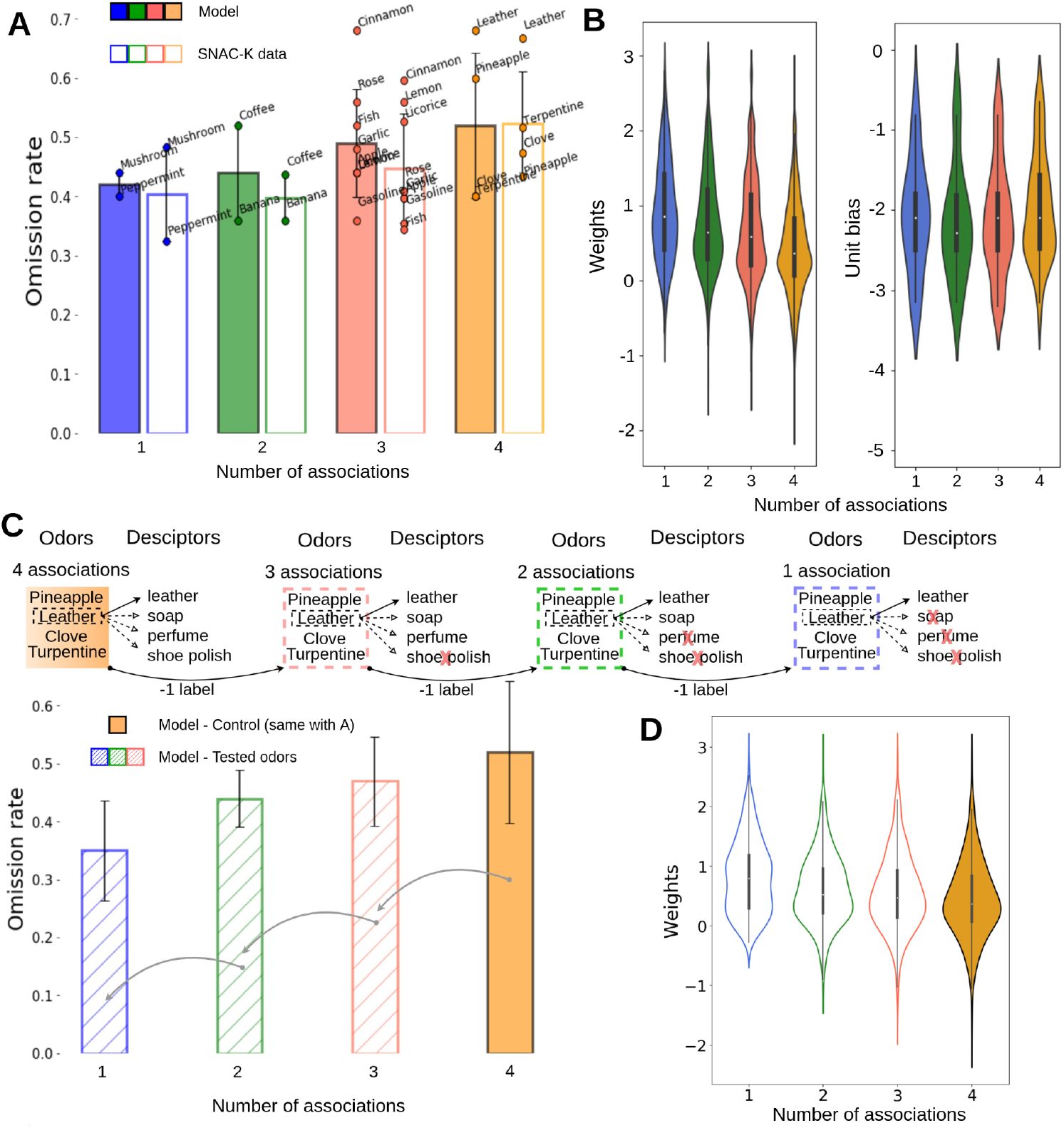
The larger number of odor-label associations, the higher likelihood of verbal omissions in model simulations. **A**) Average omission rates reported in the language network model (bars with solid fill) and behavioral SNAC-K data (bars without inner fill). The bar diagrams reveal progressive increase in the omission rates with the growing number of odor-name associations (four odor categories) established in the prior paired-associate learning paradigm. The average omission rate per odor category represents the ratio of the omissions produced by odors in the given category (the number of their word label associations) across 25 simulations. SDs represent the standard deviation of the mean across all the odors within each category. **B**) Distribution of the synaptic weights for the between-network connectivity (left) corresponding to the results in A. Weights are noticeably weaker for items which participate in multiple associations (e.g. one vs. four association). Distribution of biases of the units in the language network representations (right). Biases remain relatively balanced with the number of associations as all the odors and language entities were equally trained. **C**) We eliminated word label associations, one at a time, from odors associated with four word-labels (e.g., leather) in the original setup corresponding to the results in A. The odor label that was removed was decided by the frequency of occurrence in the SNAC-K data (responses from participants) starting from the least frequent descriptor. Here, we show an example of the odor “Leather” (top), which is initially paired with four odor descriptors in the control case. In the first iteration, after removing the odor label “shoe polish”, which is the least frequent descriptor, we re-trained the associations and evaluated the omission rates for the subset of odors with modified word label associations. We repeated this process until only one word label remained to be associated. The bar diagram (bars with diagonal lines, bottom) shows the decreasing mean omission rates in the language network subject to stepwise reduction of odor-label associations (bar filled in solid orange represents the original omission rates for odors mapped to four word labels each, the same as in A). Mean omission rates represent the mean of the averaged omissions produced by all the odors in each category across 25 simulations. SDs represent the standard deviations of the means across the odors in each category. **D**) Distribution of the between-network weights for all the odors and their associations for the simulated task shown in B.

As briefly discussed in the introduction, pairwise similarities between odor percept representations stored in the odor network and analogous similarities in the language network could also contribute to the model’s varying performance in odor recognition and naming tasks across different odors. So to examine whether our original framework for mechanistically explaining the distribution of verbal omission rates is robust, and that the observed effect is not merely driven by the similarity dependencies of some individual odors, we performed manipulations to odor-label pair bindings and evaluated their impact on the omission rates. In particular, we selected odors with the highest number of associations (e.g., four associations were present in leather, pineapple, turpentine and clove) and in a series of simulations we gradually removed their word label bindings, one at a time, to form three, two and one associations, respectively (Fig. 3C, top). It is important to mention that for each of those simulations we retrained the model (Bayesian-Hebbian paired-associate learning) to account for the modified odor-label associative mapping. The stepwise removal of word label bindings for every selected odor progressed starting from its so-called low-frequency descriptors, i.e. descriptors less frequently used by the SNAC-K participants in response to the given odor stimulus (e.g., for the odor “Leather” the least frequent descriptor was the word label “shoe polish”, which was removed at the first step of the process, Fig. 3C top). In line with our original hypothesis, we expected to observe a gradual decrease of the omission rates for those selected odors as the number of their word label associations were stepwise reduced. We observed indeed that the average trend for these “manipulated” odor memories (Fig. 3C, bottom) reflected the same overall effect, though with some odor dependent fluctuations, as reported for all the odors in their original configuration (c.f. Fig. 3A), which further corroborated our hypothesis.

What could be neural underpinnings of this progressive increase of the omission rates for odors with the higher number of associations reported in the model simulations, and what are key manifestations of paired-associate learning? To address these questions we analyzed the network elements that were subject to changes due to Bayesian-Hebbian plasticity (Chrysanthidis et al., 2022; Sandberg et al., 2002). Since the memory task involved pairing items (e.g., odors) with other items (e.g., word labels), we focused on analyzing the changes, if any, between the paired associative coupled representations. Initially, we performed analysis on the distribution of synaptic weights between the networks (between-network connectivity as indicated with red dotted lines in Fig. 2A) for the four categories of odors characterized by the number of matching word labels associated in the preceding learning process (Fig. 3B, left). In addition, we also examined the distribution of the corresponding neural biases (biases reflect changes in intrinsic excitability dynamics of units; see Methods - Plasticity learning rule) attributed to active units representing odor percept memories stored in the odor network (Fig. 3B, right). We observed that in the face of comparable biases, enforced by the balanced odor-label associative learning process, the differences in the average synaptic weights of the between-network connections for the different odor categories account for the trend in the omission rates shown in Fig. 3A. In other words, the between-network weights for the odors that were described by many word labels, thus forming many associations (one-odor-to-many-labels mapping), were weaker than for odors with fewer associations (Fig. 3B, left). In fact, we found an analogous trend in between-network weights of the model trained with modified odor-word bindings (Fig. 3D), i.e. associative connectivity became stronger when we gradually removed word labels coupled with specific odors (progressively shifting from one-to-many to one-to-few mapping). The synaptic weakening effect resulting in a weaker contribution of the stimulated odor network to the activity in the language network and, in consequence, in the higher likelihood of an omission (subthreshold response of the language network) is due to the nature of Bayesian-Hebbian learning with Bayesian Confidence Propagation Neural Network (BCPNN) learning rule. It flexibly adjusts weights over estimated presynaptic (Bayesian-prior) as well as postsynaptic (Bayesian-posterior) activity (Sandberg et al., 2002). In essence, the formation of a new association with a given odor induces synaptic weakening of all of its other associations (a form of synaptic weight normalization; Sandberg et al., 2002; Chrysanthidis et al., 2022).

### Effects of the representational overlap on omissions

The distributed representations of odor percepts and word labels, independently encoded and stored in the odor and language networks, respectively, were derived to reflect relevant similarity relationships in the external data (see *Methods - Odor percept and label representations derived from data*). However, the collective impact of the overall level of pairwise similarities between odor percept representations on the omission statistics was an open question. Since the representational similarities, parameterised in terms of representational overlap in the model (the average number of active units shared by representations of different odor percepts, see *Methods - Odor percept and label representations derived from data*), constitute an important model parameter, we globally scaled them down and re-simulated the entire free odor naming task in the model. In the light of experimental evidence suggesting that perceptual similarity between odors affect behavioral performance in both odor discrimination and identification tasks (Stevenson, 2001; Raj et al. 2023), we expected that less similarity would result in the lower omission rates in the model. Indeed, by reducing the representational overlap in both networks (odor network: mean=2.97, SD=1.06 in control vs. mean=1.82, SD=0.74 in low overlap scenario; language network: mean=3.38 SD=1.67 in control vs. mean=2.40, SD=1.04 in low overlap scenario; see *Methods - Odor percept and label representations derived from data*), the model produced considerably less verbal omissions (Fig. 4). The lowered representational overlap (reduced similarity, more distinct representations) resulted in the considerable drop in the omission rates (Fig. 4). The major reason was that more distinct representations resulted in less pattern rivalry (competition mediated by the winner-take-all circuits across the network’s hypercolumns) making it easier for the model to activate odor and then subsequently word label object in response to a partial cue (odor stimulus, see *Methods - Partial pattern activation during odor naming task in the model*). Importantly, regardless of the overlap effect (reducing the overall omissions), the omission rates exhibited the same growing tendency for odors with the higher number of word label associations as in the control case. That implies the original effect of odor-language decoupling with the increased number of associations (one-to-many mapping) was preserved even when altering the overlap between the model representations. The two effects were additive in nature.

**Figure 4:**
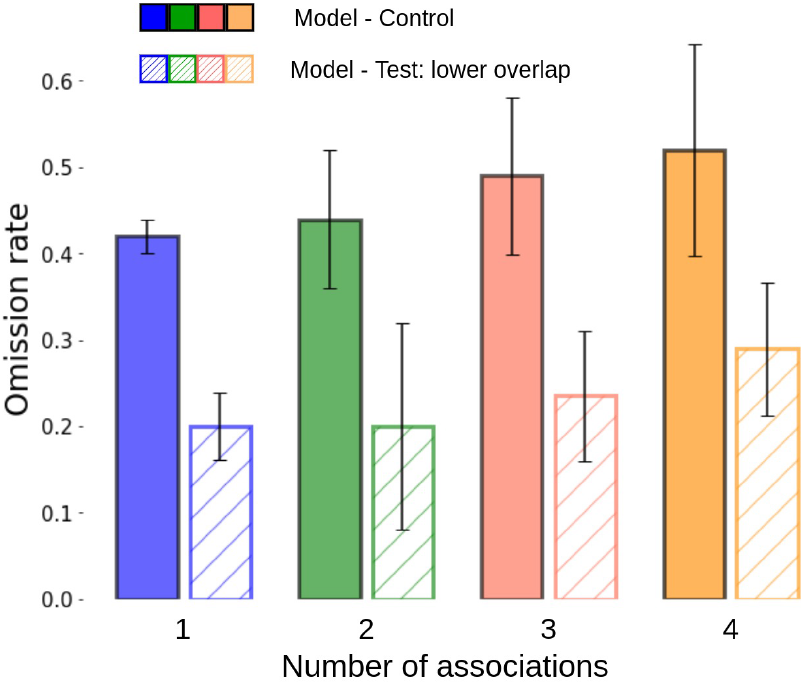
Low representational overlap in the odor and language network reduced the overall omission rates but preserved the dependency on the number of odor-label associations (across the four odor percept categories). Although the average omission rates in the language network dropped in the configuration with the low representational overlap (bars with diagonal lines) relative to the control scenario (bars with solid fill; the same as in Fig. 3A), the effect of the progressive loss of language information (i.e., decoupling) over the number of odor-name associations, reported earlier in the control scenario, was preserved. The average omission rates represent the means of the omission rates averaged over odors within each category (the number of associations to word labels) in 25 simulations and the error bars represent the corresponding standard deviations (of the means within each category).

### Effects of bi-directional coupling between the networks on omissions

In the model we assumed that odor naming builds on reciprocal interactions between the two networks. We also demonstrated the effect of the distribution of the between-network synaptic weights on the omission rates. It was not clear, however, if this effect was mediated solely by the feedforward odor-to-language activation, or also by bi-directional interactions. To gain more insights, we evaluated the consequences of disrupting the bi-directional flow of information in relation to omission rates and the odor-language decoupling phenomenon discussed earlier. In particular, we removed the language-to-odor network connectivity (Fig. 5A, control scenario in the left panel, test scenario in the right panel) and observed an overall increase in the omission rates (control: mean= 46.1 %, SD = 7.76 % vs test: mean = 53 %, SD = 10.7 % across 16 odor objects in 25 simulations; two-sample t-test: p<0.05, n=16). This confirmed our expectation that odor naming relies not only on the feedforward activation of the language network by the cued odor network but also on the reciprocal interactions between the networks. To better understand these effects, we examined individual cases of cued odor items and how the evoked network activities corresponding to the stimulated odor percept and the word labels interacted during the odor-naming recall block. We observed that the initial network response was weak, resulting in either subthreshold activity or a brief unstable (transient) activation for a limited number of timesteps. This weak odor network activity initiated a signal transmission in the language network, which in turn, through an excitatory feedback signal, amplified the odor items to maintain activity and eventually, as a result of reciprocal interactions, facilitated crossing the recall threshold in the language network (Fig. 5A, left; an example of how a closed-loop reinforcing network interaction benefits recall by producing robust and stable activations leading to a successful word label retrieval). After removing the language-to-odor network connectivity, there is no reverberation between the two networks to cause a more robust/stable odor percept activation and in turn to help the language network to converge to the corresponding label (Fig. 5A, right).

**Figure 5:**
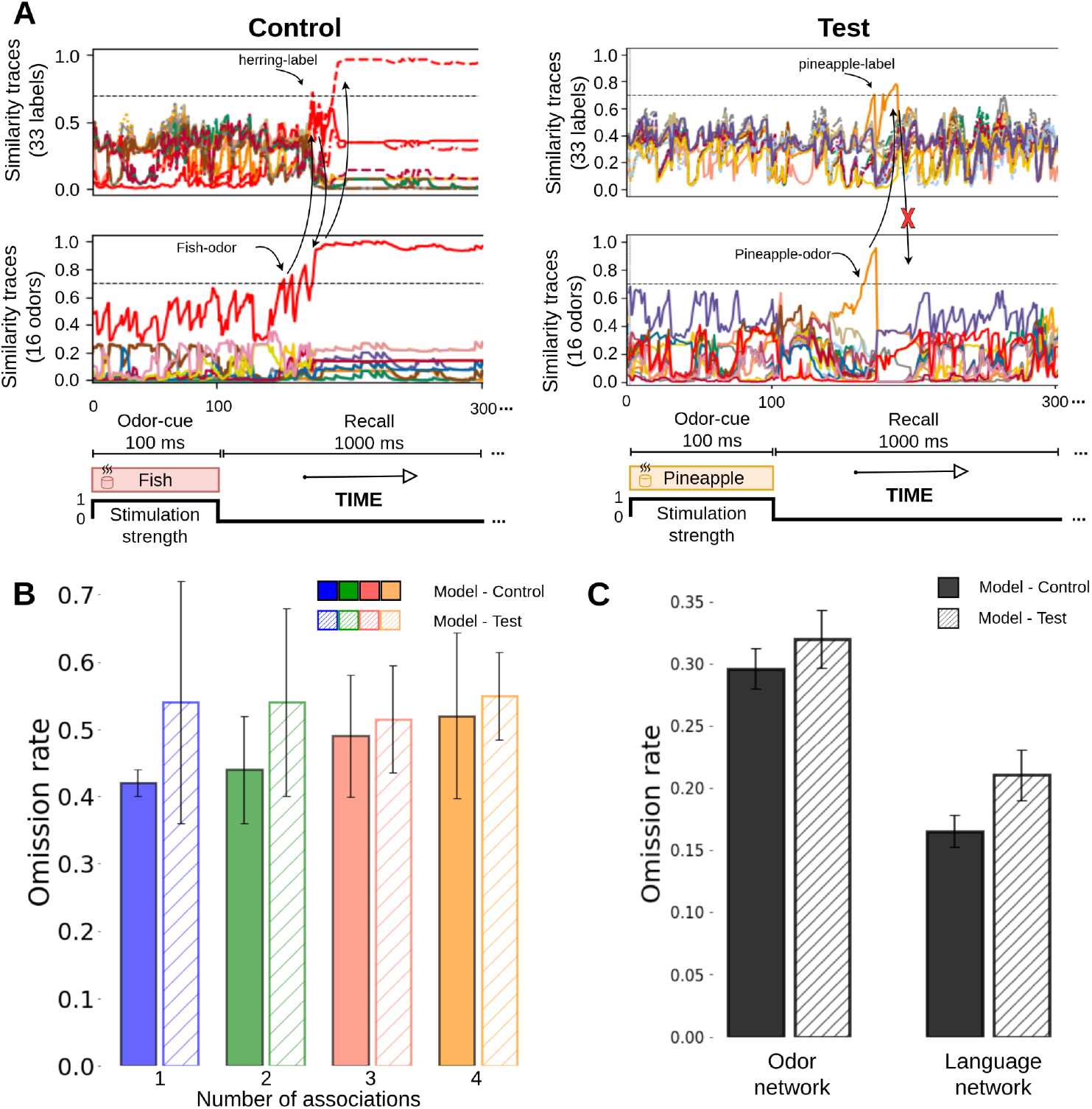
The effect of the between-network connectivity on the odor-language network interactions and odor naming performance. **A**) Language to odor projections benefit the two network interactions by amplifying the incoming signal from the odor network and spreading the activation back to the odor representations. An initial weak response of the odor network due to a partial cue can be boosted by the backward projections from the language network leading to activations of the odor percept and its label (left). However, when removing the language-to-odor connections, in some cases, the signal from the language network does not propagate back to the odor network leading to less stable attractor dynamics, manifested in either subthreshold activity or transient activation, and higher omission rates. We count as activations the convergence of attractor dynamics to a specific pattern for at least 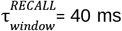 (see *Methods - Pattern activation in odor and language network*). **B**) Average omission rates reported in the language network model per odor category (the number of the corresponding word label associations) for the Control (bars with solid fill) and Test scenario (bars with diagonal lines). The average omission rate per odor category represents the ratio of the omissions produced by odors in the given category across 25 simulations. SDs represent the variability of averages across the odors within each category. **C**) Classification of the normalized omission rates to odor (perceptual) vs. language omission, for the control and test scenario.

A more systematic analysis of the consequences of removing feedback language-to-odor-network projections revealed that the previously observed trend of the increasing omission rate with the number of odor-label associations was no longer so clear (Fig. 5B). In fact, the more pronounced increase in the omission rates for the odors that follow one-to-few mapping was somewhat expected considering that their language-to-odor binding was stronger compared to the odors having one-to-many mappings (cf. Fig. 3B). So, having eliminated these feedback connections with stronger weights we could expect the relatively larger change in the omission rates for the corresponding odors. In addition, we analyzed individual simulation trials and categorized all omissions as originating in either the odor or language network. Language omission is related to inactivity of the language network when the odor network is active, whereas odor omission is caused when both networks remain silent. We first observed that lack of activation of the odor network in response to an odor cue is a more common reason behind verbal omissions than the language network deficiency (Fig. 5C). We also found that removal of the language-to-odor connections impacts both the perceptual and language omissions by increasing their rates suggesting that the reinforcing signal from the language network benefits both “odor identification” and “verbalization”.

## Discussion

In uncued odor naming tests, blank responses, omissions, are observed at an extraordinarily high frequency (Hörberg et al., 2024), even when healthy participants are smelling odors that are familiar to them (see Olofsson and Gottfried, 2015, for review). This prompted us to investigate network-level mechanisms underlying associative loss of functional connection between odors and names. In this study, our primary goal was to test a computational hypothesis that associative learning between odor and language network resources resulting in non-homogeneous distributions of synaptic associative weights could account for verbal omissions. To this end, we developed a computational model consisting of two reciprocally connected and interacting recurrent (attractor memory) networks storing long-term odor percept and word label representations, respectively. The model outcomes suggest that odor percepts that were previously associated with multiple word labels (one-to-many mapping) are relatively more susceptible to omissions, due to a weaker odor-label synaptic coupling than odors with more selective (one-to-few) label pairings. Recalling odor names evoked by an odor-perceptual experience constitutes a complex cognitive phenomenon and probes reciprocal interactions between olfactory memory and the brain’s resources underlying our capacity to use language. Despite previous neuroimaging studies on neural correlates of odor identification and a vast body of behavioral data, our insights into network-level mechanisms and into the role of synaptic plasticity in shaping the associative power of odor memories is limited, particularly in relation to odor names. Our results support a framework where olfactory and language networks interact by reciprocal connections, and where a diffuse odor-to-label mapping disrupts the activation of a word representation for verbal report. Our simulations improved our computational understanding of odor and language interactions and offered mechanistic insights into synaptic and network-level effects hypothesized to underlie the distribution of omissions observed in the experimental data. More broadly, our study contributes to our understanding of how distributed memory networks interact, offering a framework for studying associative recall processes beyond olfaction.

### Related behavioral results, and memory decoupling over longer time scales

Behavioral experiments on episodic memory using visual stimuli reported that one-to-many mapping leads to decoupling of the previously paired items (Opitz, 2010; Smith and Manzano, 2010; Smith and Handy, 2014). These studies suggest that context-specific memory traces transform into gist representations while additional memory details are progressively lost. Individual traces of item memory representations remain intact but fail to retrieve their associated representation. In fact, simple language vocabulary learning implies that learners encode words in several different contexts, which leads to a definition-like meaning of the studied word resulting in a decoupling with the associated encoded specific sentences (Beheydt, 1987; Bolger et al., 2008).

Decoupling between memories is typically described as a process that occurs over time. Full recollection of events is likely supported by multiple independent, parallel, interacting neural structures and processes since various parts of the medial temporal lobes, prefrontal cortex and parts of the parietal cortex all contribute to episodic memory retrieval including information about both where and when an event occurred (Gilboa, 2004; Diana et al., 2007; Watrous et al., 2013). A related classic idea on association degradation of memories (akin to representational decoupling) is the view that it is in fact an emergent outcome of systems consolidation. Sleep-dependent consolidation in particular has been linked to advancing the extraction of gist information (Payne et al., 2009; Friedrich et al., 2015). We acknowledge that consolidation leads to associative decoupling, yet our model proposes an additional more immediate neural and synaptic contribution to decoupling that does not require slow multi-area systems consolidation. In our work, associating odors with a variety of odor labels (one-to-many mapping) leads to decoupling of the associative memories. Our results are in line with several behavioral experiments proposing that associations of items with additional stimuli even in rapid succession is the key mechanism advancing memory forgetting (Opitz, 2010; Smith and Manzano, 2010; Smith and Handy, 2014).

Although we simulate a specific paired-associate odor naming task, the proposed plasticity mechanistic explanation that leads to odor-language decoupling may be generalized to support memory forgetting in more complex scenarios in which other mechanisms may synergistically interact. After all, our hypothesis does not exclude other seemingly coexisting phenomena and mechanisms supporting memory retrieval that may facilitate decoupling over time, e.g., reconsolidation or systems consolidation because of sleep or aging (Friedrich et al., 2015).

### Language-related vs perceptual omissions

The relative contributions of perceptual deficiencies and difficulties in odor label retrieval in odor naming failure are not well understood (Olofsson and Gottfried, 2015). In a recent study, Lindroos et al. (2022) examined different correlates of omissions in an olfactory experiment. They assessed the perceived pleasantness, familiarity, intensity, and edibility of odors and examined how odor identification performance was associated with these variables. A key finding linked odor intensity with enhanced task performance, that is, high-intensity odors were easier to identify, suggesting that high intensity ratings may improve recognition of the perceptual level. Understanding factors underlying the omissions and their systems level correlates is essential for accurately interpreting task outcomes and improving the reliability of odor assessments.

In this work, we computationally investigated two categories of the origins of omissions: those arising from olfactory perceptual issues (attributed in the model to the odor network) and those broadly related to the language system (irresponsive language network despite the activation of the odor network in the model). Our simulations suggested that the high omission rates in a free odor naming task paradigm may stem primarily from deficiencies in the perceptual odor domain. This issue appears to be driven by the significant overlap among different odors increasing the competition in the network. Although the ability to view a banana and conjure up the word ‘banana’ comes easily and rapidly, the corresponding ability to smell a banana and conjure up the word ‘banana’ can be extremely hard (Olofsson and Gottfried, 2015), possibly due to the strong similarities at the perceptual level. When we lower the total overlap between representations we observe that the odor naming improves. We further noticed that by testing orthogonal odor representations (which do not capture the inter-relationships between odors with high similarity) the resulting perceptual omissions become even more reduced, boosting our findings that overlap may play a critical role in odor naming omissions. In conclusion, memory representations with high overlap with other patterns may result in higher omission rates compared to more distinct representations.

### Related abstract models of familiarity and recollection

Perceptual or abstract single-trace, dual-process computational models, based on signal detection theory, used to explain memory recall (Wixted, 2007; Greve et al., 2010), often aim to explain traditional Remember/Know behavioral studies. Participants in such studies are instructed to give a “Know” response if the stimulus presented in the test phase is known/recognized or familiar without any further accompanying memories regarding the stimulus. Conversely, “Remember” judgments are to be provided if the stimulus is recognized along with some recollection of specific information. This results in a strict criterion for recollection, as it is possible for a subject to successfully recognize an item but fail to retrieve the source information (Ryals et al., 2013). It is worth mentioning that these computational models may explain memory recall but the potential loss of additional information is only implied as it does not have its own independent representation. However, our dual-network model offers distinct representations (here: odor and language networks) that help us pinpoint the sources of loss of information. Mandler (1979, 1980) and Atkinson and Juola (1973) treat familiarity as an activation of preexisting memory representations. Our results are compatible with this notion because our model proposes to treat item-only (odor) activations corresponding to verbal omissions as “Know” judgments, while those accompanied by the activation of language representations best correspond to a “Remember” judgment.

### Biological network plausibility

#### Modular cortical architecture

Our model with populations of neural units organized in columnar modules aligns with data regarding cortical functional architecture (Mountcastle, 1997; Hubel and Wiesel, 1977; Buxhoeveden and Casanova, 2002; Favorov and Kelly, 1994; Rockland and Ichinohe, 2004; Yoshimura et al., 2005). These functional columns share high degree of recurrent connectivity (Thomson et al., 2002; Yoshimura and Callaway, 2005), which also facilitates the formation of larger modules - subnetworks such as cell assemblies (Stettler et al., 2002; Binzegger et al., 2009; Muir et al., 2011; Eyal et al., 2018). Spanning an area with diameter about 500 μm (Mountcastle, 1997), these columns are spatially distributed in the cortex. Their functional minicolumns (populations of tightly recurrently connected excitatory neurons within the superficial layer of columnar modules, parsimoniously modeled in this work as units) intermingle with similar modules of other cell assemblies (Perin et al., 2011). Our model incorporates a local soft winner-take-all architecture within each hypercolumn integrating the functionality of fast-spiking local inhibitory interneurons such as basket cells that provide the primary source of feedback inhibition, which leads to competition similar to that of a local soft winner-take-all module.

#### A simplified framework for odor-language system interaction (two-network system architecture)

The literature supports an object-based approach to odor-source naming. Objects (e.g., the smell of popcorn) constitute building blocks of perception and provide the input to language subsystems for source naming. The striking mechanistic similarities between human and rodent data (Wilson and Stevenson, 2006) and between vision, audition, and olfaction (Gottfried, 2010) lead us to believe that odors are universally encoded as discrete objects. This object-based encoding aligns well with attractor dynamics, which play a crucial role in odor (object) recognition (Kay et al., 1996). Naturally, in the olfactory system, associative representations in the piriform cortex are densely and reciprocally integrated with limbic and paralimbic areas in the medial temporal lobe and basal forebrain (Zahnert et al., 2024; East et al., 2021). In humans, such pathways offer opportunities for endowing olfactory object information with limbic associative systems. In a free odor naming paradigm the feedforward projections from the piriform cortex mediate relatively coarse odor object information directly to the language network system via the orbitofrontal cortex and temporal pole (Olofsson and Gottfried, 2015). The interaction between odor and language information is complex. In our computational approach we relied on a parsimonious model for establishing memory associations between olfactory perception objects, i.e. odor percepts with correlates predominantly in the piriform cortex, and odor labels generated broadly by the language system. While these brain systems, language in particular, comprise distributed network of networks and their inter-connections are also mediated or influenced to some extent by other networks involved in the recollection process (e.g. orbitofrontal cortex, temporal pole; Olofsson and Gottfried, 2015), we intentionally reduced the model to two long-term-memory-like networks with reciprocal connectivity so that their interactions are in the focus. Consequently, we considered these two subsystems to store pre-encoded (consolidated) representations of odor percepts and odor labels, and assumed that paired-associate learning linking odor percepts with their names also takes place over a long time span (past episodic experiences).

#### Bayesian-Hebbian plasticity and other Hebbian-like learning rules

Our model employs a Bayesian-Hebbian associative learning rule, yet there are other associative Hebbian-like learning rules also used in computational studies, e.g. classical Hebbian (Herz et al., 1989) or covariance learning rule. We cannot exclude that an alternative plasticity learning rule may mechanistically explain odor-naming decoupling and produce comparable effects. However, the key strength of the Bayesian-Hebbian learning rule embodied by BCPNN lies in the normalization and weight update over estimated presynaptic (Bayesian-prior) as well as postsynaptic (Bayesian-posterior) activity (Sandberg et al., 2002). This encapsulates multiple synaptic and neural processes at varying biologically relevant time scales, and balances the formation of new and gradual forgetting of older correlated associations (Chrysanthidis et al., 2022), unlike the classical Hebbian-like learning rule. In fact, in our recent study (Chrysanthidis et al., 2022) we have shown that a detailed spiking Hebbian learning rule in the form of a conventional spike-time dependent plasticity (STDP), unlike BCPNN, does not explain this memory decoupling effect between associated representations.

#### Future work

In future work we intend to explore the model’s ability to predict correct and incorrect odor label responses, e.g. labeling a garlic odor as “fruit”. While our current model generates both correct and incorrect responses, aligning model statistics with experimental findings remains a key objective which will initiate further exploration on other related network-level mechanisms. To achieve this, additional odor information, such as familiarity, edibility and intensity, may have to be integrated at the individual odor level (cf. Hörberg et al. 2024). This integration would allow model responses to drift from incorrect to correct, and vice versa, depending on the levels of these parameters.

It should be noted that we observed significant variability in the model among individual odors despite convincing decoupling effects at the odor group level. In other words, individual omission rates for specific odors often deviated considerably from the average omission rates of the odor group. For instance, cinnamon, forming three associations, exhibited significantly higher omission rates compared to the mean average (Fig. 3A). We did not find this variability surprising, as prior analysis of omissions in SNAC-K data already revealed that omission rates for some odors deviated from the average omission rates of the odor group (Fig. 1; and see Hörberg et al. 2024). Our future investigations will seek to elucidate how unique odor characteristics, such as varying levels of intensity or familiarity, may contribute to these individual differences in variability. In this study our emphasis was on summative effects underlying general trends in verbal omissions in a free odor naming task paradigm and we only accounted for perceptual similarity between odors without considering other individual odor features in the generation of the embedded odor memory representations.

## Methods

### Computational model

#### Network model architecture

The dual-network model architecture, inspired by our previous work on item-context episodic memory binding (Chrysanthidis et al., 2022; Chrysanthidis et al., 2024), rests on two recurrently connected memory networks (Fig. 2B, within-network connectivity), which are also associatively connected (Fig. 2B, between-network connectivity). One network accounts for an olfactory memory system that stores odor percepts (odor network), and the other network corresponds to a language reservoir of linguistic labels at the given level of abstraction (language network). We utilized a less detailed non-spiking implementation with population rate based coding units, with both networks having the same modular architecture, inspired by the columnar organization of the mammalian neocortex (Mountcastle, 1957, 1997; Hubel and Wiesel, 1962; Douglas and Martin, 2004; Bastos et al., 2012). In particular, each associative recurrently connected memory network is composed of units, which encapsulate the superficial L2/3 portion of cortical minicolumns (MCs), organized into modules, referred to as hypercolumns (HCs). As mentioned, each minicolumnar unit represents a local population of units that are recurrently connected to reflect the local excitation within a functional cortical module. The units locally compete within the shared hypercolumn module, defined as the extent of locally shared lateral inhibition, by means of a soft winner-take-all operation via divisive normalization of activity. Thus, the network is composed of several identical modules, every one of which in turn comprises 15 minicolumn units. Importantly, the dense long-range connectivity between HCs links (Thomson et al., 2002; Yoshimura and Callaway, 2005) coactive modules into larger cell assemblies forming attractor memories (Stettler et al., 2002; Binzegger et al., 2009; Muir et al., 2011; Eyal et al., 2018). The basis for their formation is Bayesian-Hebbian-like synaptic plasticity, modeled as the BCPNN learning rule (see *Plasticity learning rule* in *Methods*).

The active units are subject to adaptation (*a*_*j*_) with the time constant *τ*_*α*_. The total input to the j-th unit (*s*_*j*_), and its output rate (*o*_*j*_) are updated according to the following equations (see Table 1 for the default parameter values):

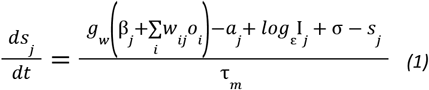

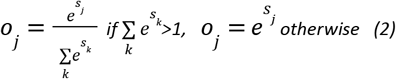

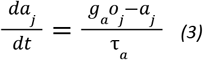

In our framework, we denote connectivity gain (*g*_*w*_), and adaptation gain (*g*_*α*_), as well as the unit time constant (*τ*_*m*_). Additionally, I_j_ represents the input to unit j, and σ signifies the amplitude of zero-mean Gaussian noise incorporated into the support of each unit per iteration. The temporal evolution of unit activation resulting from synaptic input, adaptation, external input, and noise is reflected in Eq. 1. The model implements a half normalization of unit activity within its hosting hypercolumn module, which reflects the divisive lateral inhibition (Eq. 2). The adaptation a_j_ is governed by an exponential decay process with the time constant of *τ*_*m*_ (Eq. 3).

#### Plasticity learning rule

We implement synaptic plasticity using the BCPNN learning rule (Lansner and Ekeberg, 1989; Wahlgren and Lansner, 2001; Sandberg et al., 2002). The BCPNN synapse continuously updates three synaptic biophysically plausible local memory traces of pre-, post- and co-activation, *p*_*i*_, *p*_*j*_ and *p*_*ij*_, respectively, from which the Bayesian bias and weights are calculated (Sandberg et al., 2002). Learning implements exponential filters, *z*, and *p*, of activation with a hierarchy of time constants, τ_*z*_, and τ_*p*_, respectively. Due to their temporal integrative nature they are referred to as synaptic (local memory) traces. ln more detail, the pre- and post-synaptic trace variables (*z*) represent presynaptic and postsynaptic activation involved in synaptic plasticity like e.g. NMDA channel opening (*z*_*i*_) and membrane depolarization due to back-propagating events (*z*_*j*_). These traces typically have relatively short time constants (*τ*_*zi*_, *τ*_*zj*_) serving as a short-term memory buffer allowing modification of connections between units that are activated within a short time window. The *z*-traces are calculated according to the following equations:

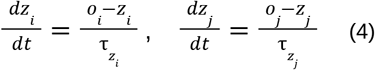

The estimated probabilities of pre-(*p*_*i*_) and post-synaptic (*p*_*j*_) activation and co-activation (*p*_*ij*_) are updated as:

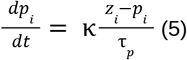

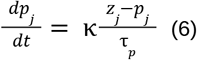

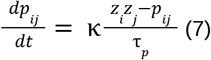

This tends to bind units that are activated together via excitatory learned weights. There is a separate longer time constant corresponding to the learning rate for the trace variables (*τ*_*p*_), and a gating parameter for learning (*κ*), as well as a gain for the neural bias (*g*_*b*_). The parameter *κ* adjusts the learning rate. Setting *κ*=0 freezes the network’s weights and biases, though in our simulations the learning rate remains constant (κ_*encoding*_=1) during encoding and κ_*recall*_=0 during recall. The connections weights (*w*_*ij*_) and biases (*β*_*j*_) are continuously computed based on these p-estimates according to the Bayesian-Hebbian learning rule (Sandberg et al., 2002) as:

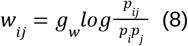

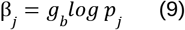

The bias term represents the log prior of the post-synaptic unit and represents the elevated baseline activity of units frequently activated by their inputs (hence it is sometimes referred to as intrinsic excitability; Fransén et al., 2006; Jung and Hoffman, 2009).The weight term accounts for the mutual information between pre- and post-synaptic units. It needs to be pointed out that our learning rule produces excitatory connections between units with correlated activation and inhibitory connections between units with anti-correlated activity in the spirit of Hebbian learning. In particular, units that are not active together in a certain time window feature low coactivation traces (*p*_*ij*_), and based on Equation 8, the final weight update produces negative weights. This negative binding assumes that the inhibition is disynaptic, mediated via local inhibitory interneurons, which may play an important role in shaping neural activity (DeFelipe et al., 2006; Krimer et al., 2005; Kelsom and Lu, 2013; Chrysanthidis et al., 2019). We do not explicitly model this disynaptic inhibitory circuit but instead use negative weights in network computations.

#### Memory pre-encoding

Encoding odors and labels: before simulating the SNAC-K free odor naming task, the odor and label patterns underwent a training process to establish well-consolidated memories. Initially, each memory pattern was encoded separately using one-shot learning, where each pattern was trained for 100 ms with Hebbian learning, employing long time constants across multiple simulations (Sandberg et al., 2002). Once this initial encoding was complete, we sampled network elements (e.g., synaptic weights) to construct the odor and label representations at their corresponding networks.

Encoding odor-label associations: to bind the label and odor representations, odors were first grouped into sets of four groups based on the number of their paired associations (see Fig. 2A). For odors that were paired with four different labels, each odor was activated sequentially with each of the four labels for 100 ms. Similarly, for odors with two associations, each odor-label pair was activated twice, maintaining a total of four activations per odor. To avoid any biased learning effects in the odor network due to varying number of label associations per odor, the stimulation protocol was balanced so that each odor was cued the same number of times. This learning protocol ensured that all odors were activated the same number of times, balancing the activation duration regardless of the number of associations. The paired cue paradigm was implemented using slow Hebbian plasticity, as before. At this stage, the between-network connectivity (Fig. 2B) was being shaped reflecting odor-label pairs (Fig. 2A) while the local within-network connectivity had been stabilized through previous training.

Finally, to simulate the SNAC-K odor naming task, both local and between-network connectivity were preloaded into the network.

#### Model stimulation with an odor perceptual cue

During the simulated task, the odor network is stimulated for 100 ms with a partial cue corresponding to a selected odor percept with the following recall phase of 1000 ms. The external cue targets a random subset of 10 units of the stimulated odor representation (that is encoded with 15 units, one in each columnar module). Prior to each partial cue, the network was reset to a default state to exclude the possibility that ongoing activity induced by previous trials influenced the network state for new stimuli.

#### Pattern activation in the odor and language network

We quantified each network’s response by calculating the cosine similarity between the distributed pattern of network activation and the memory representations of all stored objects. This approach allowed us to track the odor percept and word label that the respective network responses converged over time. Once the resulting convergent state was sufficiently similar to one the stored memory states (i.e. the recall detection threshold *r*_*th*_ was exceeded for at least a certain duration 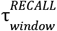, see Table 1), that memory activation, i.e. odor percept for the olfactory network or word label for the language network, was considered as a perceptual or verbal output, respectively. The subthreshold activation of the language network within the 1-s-long recall block, i.e. no legitimate output, was considered as a verbal omission.

### Behavioral data, analysis and modelling perspective

#### Odor naming behavioral task

The data that we used to compare our model output against was collected from a Swedish National Study on Aging and Care in Kungsholmen (SNAC-K), where 2569 subjects underwent an odor naming test of 16 odors (see Larsson et al., 2016 for a detailed description). For the analysis in this study, participants diagnosed with dementia (n = 60), Parkinson’s disease (n = 20), Schizophrenia (n = 9) and developmental disorders (n = 1) were excluded. The effective sample thus consisted of 2479 participants with the mean age of 71.80 years (61.3% women). The following odors were included: apple, banana, cinnamon, cloves, coffee, fish, garlic, leather, lemon, licorice, mushroom, peppermint, petrol, pineapple, rose, and turpentine. The SNAC-K odor identification test was a slightly modified version of the Sniffin’ Sticks identification test (Hummel et al. 1997). After presenting each individual odor, the SNAC-K participants were asked to name the odor without any further cues (odor naming in a free identification task). Participants’ responses were categorized as hits, mistakes, and omissions. If the answer was not correct (mistake or blank response to a test cue), participants were provided with four alternatives, three distractors and one target odor-name (cued identification) displayed simultaneously on the screen.

#### Odor similarity behavioral task

The similarity of the odors in SNAC-K were collected in a separate behavioral experiment by Lindroos et al., (2022). Thirtyseven participants (age range 19–61 years, 20 females and 17 males) were provided with all the possible pairs between the 16 odor labels used in SNAC-K, presented in a random order, and rated their pairwise similarity on a visual analog scale from 0 (no similarity) to a 100 (identical). All participants rated each pair twice, where the order of the presented odors was reversed: first odor A was followed by odor B and in the second assessment odor B was followed by odor A. In total 240 pairs were rated by each participant (see Lindroos et al. (2022) for a description of the cohort, and additional information on the task conditions and instructions). The average similarity of each pair were then calculated and stored in a matrix that were then used to produce odor representations for our model.

#### Odor percept and label representations derived from data

In studies of human capacity of odor naming it is vital to account for perceptual similarities between odor stimuli as well as semantic relationships between linguistic labels. In our simulations we incorporated these factors in the network representations of both odor and word label objects stored as attractor memories in the model. This was done by creating attractors with a relative overlap of the units resembling the similarity of the odors and words, respectively. Despite their abstract nature, i.e. binary vector representations of active and inactive units in the network, they can be made more or less similar using a suitable metric such as scalar product that quantifies the level of representational overlap.

Sparse distributed representations of odor percepts and labels, embedded in the memory model, were created by matching the pairwise similarities between, respectively, odor percepts and Word2Vec odor label embeddings (Fig. 6, left). Word2Vec assesses the semantic similarity between words based on contexts within a large corpus of text (Mikolov et al., 2013), here: the Swedish Blog corpus, see Östling & Wirén 2013. The mapping was created using our own implementation of the Greedy Max Cut method (GMC, Rohde, 2002), in such a way that the number of overlapping minicolumns of the attractors should reflect the similarity measurement between the odors (Fig. 6, left, middle). This was done by looping over hypercolumns and patterns one-by-one: for each hypercolumn and pattern the minicolumn which minimized the global mean difference between the pairwise Hamming distances (Norouzi et al., 2021) of the attractors and the distance matrix of the odors, was selected (Fig. 6, right). This was then repeated over and over until the error was minimized (Fig. 6, right, differences between semantic similarities followed a similar trend). The python code implementing this transformation is available online (see Data availability).

**Fig. 6:**
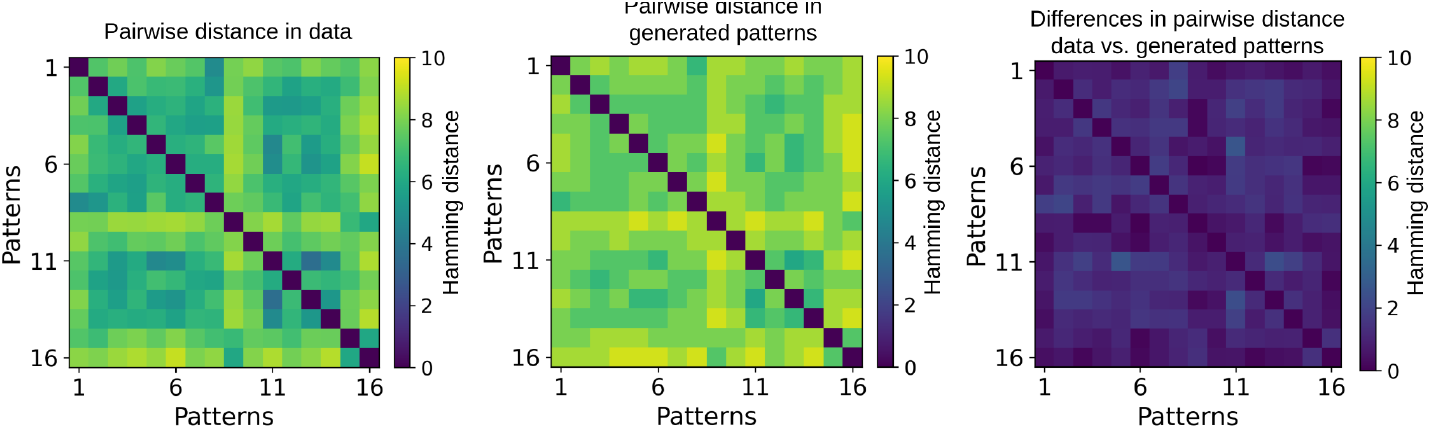
Pairwise distances between odor percepts in the experimental data, left. Pairwise distances of the generated patterns using the GMC algorithm, middle. The resulting patterns were used as input to the model, middle. Absolute differences in pairwise distances between odor percepts in the data vs. generated patterns, right. The generated input patterns closely approximated the data, as the pairwise distance differences between odor percepts in the data and the generated patterns were minimal.

Each odor and language representation was composed of 15 minicolumns (Table 1), with some of them shared with other odor and language representations depending on their respective perceptual and semantic similarities (overlap in odor network: mean=2.97, SD=1.06; mean reflects the average number of shared units across all the odors and SD defines the variability/width of the distribution; overlap in the language network: mean=3.38 SD=1.67). Eventually, the produced sets of sparse activity patterns were encoded, before actually simulating the experimental trials (Memory test), with long-term Bayesian-Habbian learning in the network representing consolidated odor and label memories.

#### Characterizing odor-label diversity

To measure the diversity of word labels used by the participants of the SNAC-K study to name each odor, we used the Simpson diversity index (Majd et al., 2018). It has been previously used to quantify naming diversity taking into account both the type and frequency of word labels per odor stimulus. For a given stimulus within a language the Simpson’s diversity index is as follows:

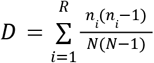

where n refers to the total number of responses using that particular name and N is the total number of responses across all names. An index of 1 indicates low label diversity - all respondents produced the same description, whereas 0 indicates high label diversity since all respondents produced different unique descriptions. For each odor stimulus we calculated the label diversity and then compared these values (see Fig. 1 and Results).

## Data availability

The BCPNN learning rule and the code that was used to run the experiments is available publicly on GitHub and can be accessed at https://github.com/Nikolaos-Chrysanthidis/Odor-naming.

In addition, the algorithm generating the odor percept and label representations using collected data can be downloaded from: https://github.com/robban80/greedy_max_cut.

## Acknowledgements

This research was supported by Vetenskapsrådet (2018-05360, 2018-07079 and 2020-00266), the Swedish e-Science Research Centre (SeRC), Digital Futures, and European Commission, Directorate-General for Communications Networks, Content and Technology (101135809) and Knut and Alice Wallenberg Foundation (2016:0229). The simulations were performed on resources provided by National Academic Infrastructure for Supercomputing in Sweden (NAISS) at the PDC Center for High Performance Computing, KTH Royal Institute of Technology.

Conflict of interest statement: The authors declare no competing financial interests

